# Quantifying cells : moderately increasing stomatal density through overexpressing *FSTOMAGEN* improved the biomass of *Arabidopsis*

**DOI:** 10.1101/2025.06.27.661899

**Authors:** Yong-yao Zhao

## Abstract

Stomata are the pores on plant surface, and these tiny pores are responsible for the flow of gas between plants and atmosphere. Currently, whether quantifying stomatal density to a moderately high level via genetical manipulation could improve *Arabidopsis* biomass remains unknown.

The Arabidopsis transgenic lines with increased stomatal density were acquired through overexpressing a gene (the homologs of *STOMAGEN, FSTOMAGEN*, the homologs in *Flaveria*). The biomass of *Arabidopsis* transgenic lines with intermediate stomatal density increased. Compared with the other lines, the biomass of the transgenic lines with excessive stomatal density exhibited no difference or decrease. There was a positive and significant correlation between biomass and relative water content. During the earlier phase of growth, the leaf growth rate and leaf area of the lines with higher stomatal density was faster and larger respectively, and there was a significant linear relationship between stomatal density and leaf growth rate and a strong linear relationship between stomatal density and leaf area, which was in agreement with the notion that more stomata indicated faster CO_2_ assimilation and then faster accumulation of biomass, although it occurs in earlier phase. This suggested that the increased biomass was attributed to the earlier phase.

Results indicated that finely tuning stomatal density could improve *Arabidopsis* biomass by genetically manipulating *FSTOMAGEN*. The functions of *STOMAGEN* homologs are extremely conserved in the evolution, thus it is promising that this genetically modified strategy could be applied in other species, which benefits the reduction of the concentration of atmosphere CO_2_ and the production.

## Introduction

Stomata are the tiny pores on the surface of plants. They are the passages for the transfer of gas (e.g. CO_2_, H_2_O, O_2_) between plants and atmosphere. Passing through stomata, a large amount of water loses from plant and spreads into atmosphere. Therefore, stomata are apparatuses which control the water status of plants. Besides, stomata also participate in the photosynthesis, since CO_2_ enters into plants through stomata. Therefore, stomata are also responsible for the fixation of carbon in plants. Because stomata are on the surface of plants, the methods for observing stomata are easy, and the morphology of stomata is very easy to distinguish. These enable stomata to become a model to study the development of cells, especially, in plants. Due to these reasons, stomata have been deeply and widely studied.

The mechanism of stomatal development has been well studied, and many genes related to stomatal development have been found (Pillitteri and Torii 2012; Zoulias et al. 2018). *EPF*family contain a series of genes which encode signal peptides(Takata et al. 2013; Hunt and Gray 2009; Hara et al. 2007; Sugano et al. 2010; Kondo et al. 2010; Hunt et al. 2010). For these genes, more focuses are put on *EPF1, EPF2* and *STOMAGEN*. The genes in EPF family *EPF1/2* negatively regulate the development of stomata, and *EPFL9* (*STOMAGEN*) positively regulates this development(Hara et al. 2007; Hunt and Gray 2009; Sugano et al. 2010; Kondo et al. 2010; Hunt et al. 2010). The *EPF1/2* mutant and overexpression lines have the uniformly changed stomatal density in multiple species(Wang et al. 2016; Hughes et al. 2017; Caine et al. 2018), and *STOMAGEN* mutant and overexpression lines also have the uniformly changed stomatal density in various species(Lu et al. 2019)(Tanaka et al. 2013; Shahbaz et al. 2025), suggesting that these genes in *EPF* family are functionally conserved during evolution. The changes in these genes could affect the stomatal density, therefore many works change stomatal conductance through genetically manipulating these genes, and thus change transpiration and photosynthetic rate(Tanaka et al. 2013; Franks et al. 2015). Therefore, because of the changed photosynthetic rate, in theory, the changes in stomatal density likely cause the consequences that plant biomass is changed in these genetical engineering processes. Increased photosynthetic rate caused by raised stomatal density likely leads to increased plant biomass. However, work shows that the overexpression of *STOMAGEN* can’t increase the biomass of *Arabidopsis*, though stomatal density of the line with overexpression of *STOMAGEN* dramatically increases(Tanaka et al. 2013). The possible reason for this increased biomass might be the overmuch stomata in the transgenic lines, since it is possible that overmuch stomata will result in the excessive loss of water in plants and consume the excessive loss of energy which participates in stomatal movement and development. Therefore, our study hypothesized that moderately increasing stomatal density improves the biomass. Recently, stomatal density modified through genetical engineering has been raised a finely tuned level, influencing the drought resistant(Nicholas G. Karavolias 2023). In this study, they found only the lines with moderately decreased stomatal density could improve the drought resistant without the accompanying of decreased stomatal conductance, carbon assimilation and thermoregulation(Karavolias et al. 2023). This study supports the notion that changing stomatal density to a moderate level might be advantage than an overmuch level. However, currently, whether increasing stomatal density to an moderate level by genetically manipulating could increase biomass remains unknown.

In this study, we examined (*FSTOMAGEN*) overexpression lines with different stomatal density in *Arabidopsis* . Our hypothesis is that stomatal density is increased to an moderate level by genetical manipulation could accompany the increased biomass of *Arabidopsis* .

## Methods

### 1 Building construct, transforming and acquiring the transgenic plants

We over-expressed the functional region of *FSTOMAGEN* (the homologs of *STOMAGEN* in *Flaveria*) into *Arabidopsis* (Zhao et al. 2022). Specific speaking, the 35S promoter was used to drive the overexpression of the functional region of *FSTOMAGEN*. The functional region of *FSTOMAGEN* was acquired through the amplification from cDNA of *F*.*rob*. The signal peptide of the gene in the lines used in this study was also amplificated from *F*.*rob*. These molecular fragments were installed in the pCAMBIA-3300 carrier through homologous recombination(Vazyme). The pCAMBIA-3300 carrier was transformed into colon bacillus. Then, the carrier was transformed into agrobacterium GV3101(VEIDI). Transgenic lines were acquired by flower-dipping method. The transgenic lines were selected with 2000-diluted Basta. T2 generation of transgenic *Arabidopsis* was performed for subsequent experiment.

### 2 Growth condition

For the experiment, at the beginning, the transgenic *Arabidopsis* lines were grown under about 100 PPFD. After 19 days, the transgenic *Arabidopsis* lines were transferred into about 300 PPFD. Then, these transgenic *Arabidopsis* lines were transferred into about 500 PPFD. During the whole growth period, sufficient water was given to avoid drought and excessive moistening in soil.

### 3 The measurement of biomass

Biomass was tested by using the weight of *Arabidospsis*. The weight was divided into two definition: fresh weight and dry weight. The *Arabidopsis* was putted into the oven to be the biomass without water, and the dry weight can be acquired. Both dry weight and fresh weight was measuring by using Electronic Balance (sartorius).

### 3 The calculation of water content

The relative water content was calculated with following equation:

(The fresh weight – The dry weight)/The fresh weight

### 4 The photoing of *Arabidopsis*

After transfer the plants to soil, waiting 7 days and then photoing plants, and the interval of the photoing was two days. The photoing was performed by Cannon EOS 1500D camera (Canon Inc., Japan)

### 5 The graphing, plotting, the statistical analysis and linear regression

The student t test (unpair, two tail) was perfomed by Graphpad prism 8 or excel. The linear regression was performed by Graphpad prism 8 or excel. The growth curve was performed by Graphpad prism 8. The model used in the growth curve was the growth model which is embedded in Graphpad prism 8. The Internal studentized residual and External studentized residual were performed by artificial intellegence, and the artifcial intellegence was name as Doubao which was developed by ByteDance. The graphing and plotting were perfomed by R language (ggplot) and graphpad prism 8.

## Results

### 1 The transgenic lines with moderately increased stomatal density showed increased rosette area and biomass

The 9 *Arabidopsis* ransgenic lines with the overexpression of the gene (*FSTOMAGEN*, the homologs in *Flaveria* of *STOMAGEN*) were acquired. Our work showed that stomatal density in these lines increased stepwise, and there was a positive relationship between stomatal density and expression level(Zhao et al. 2022). Besides, the work from others also indicates that the expression level of *STOMAGEN* among the different lines with the overexpression is positively correlated with stomatal density(Sugano et al. 2010).

Markedly, the differences in the total leaf area between these transgenic lines were observed (Fig 1A). Compared with other lines, two lines (11 and 12 line) had apparently increased total leaf area (Fig 1A). This encouraged us to test the biomass of these transgenic *Arabidopsis* lines. In consistent with total leaf area, there was a variance in biomass between these lines. The changed trend in fresh and dry biomass, accompanied by the stepwise increase in stomatal density, was first rise and then decrease (Fig 1 C, D). Compared with the line with lowest stomatal density (8 line), the medians of fresh and dry biomass in 11 and 12 lines were higher (Fig 1 C, D). Particularly, compared with 8 line, the dry and fresh biomass of 12 line significantly differed (Supplementary file). The difference in dry biomass between 11 and 8 lines was in the margin of statistic test (Supplementary file). The stomatal densitys in 30 lines and 9 lines were higher among all these lines, but the biomass in 30 lines was significantly lower than that in 8 line and there was no significant difference in biomass between 9 lines and 8 lines (Supplementary file).

**Figure 1.**
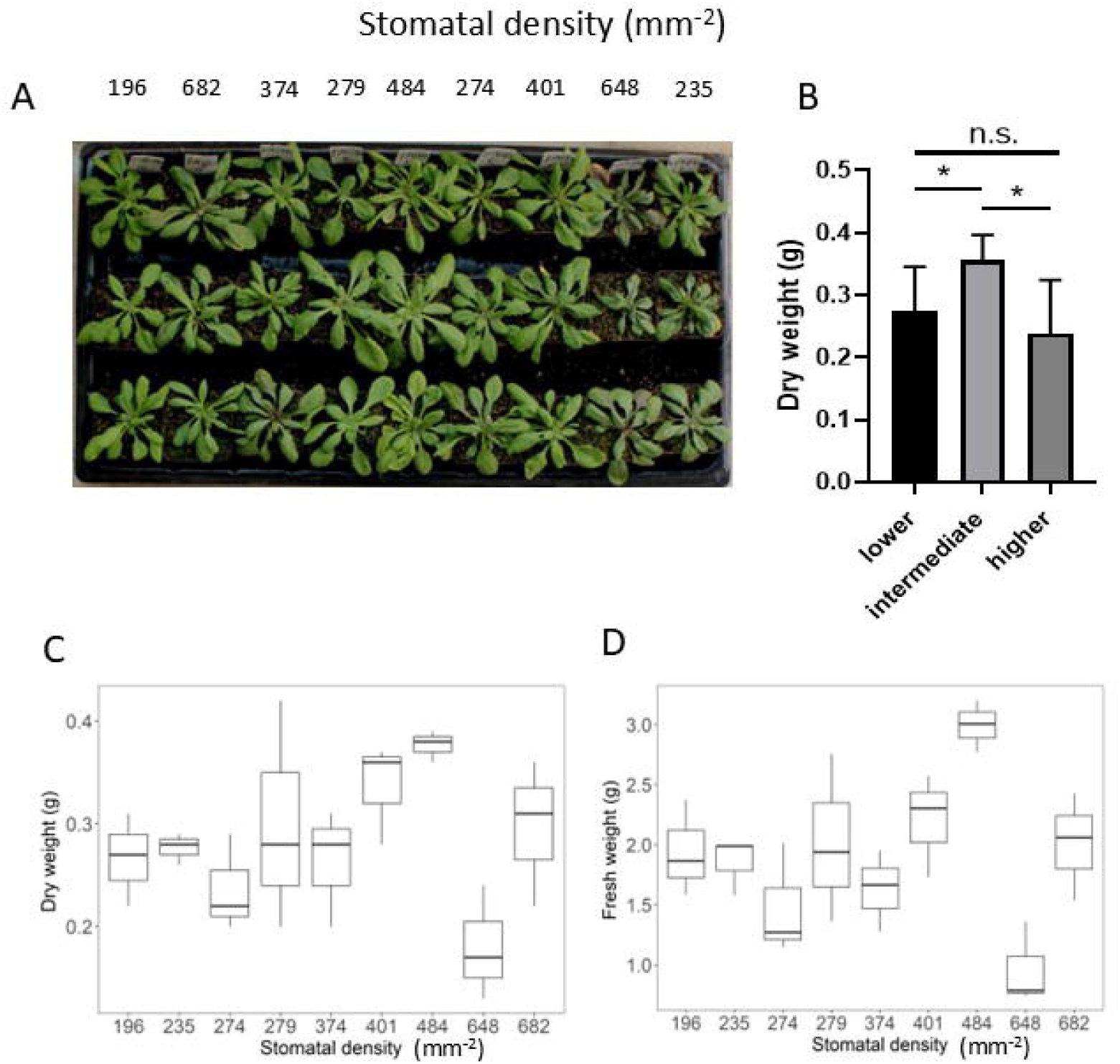
The changes in biomass between different *Arabidopsis* lines with the *FSTOMAGEN* overexpression. A, The changes in rosette morphology between different *Arabidopsis* lines with different stomatal density, the stomatal density is shown above the photos of *Arabidopsis* . B, The changes in dry weight between three levels of stomatal density. Because there was an apparent trend in the changes in dry weight (Fig C), we classified the lines into three groups according to their stomatal density. C, The changes in dry weight between different *Arabidopsis* lines with different stomatal density. D, The changes in fresh weight between different *Arabidopsis* lines with different stomatal density. The different stomatal density was achieved by the overexpression of *FSTOMAGEN*, and the different stomatal density was achieved by the different expression level of *FSTOMAGEN* in the overexpression. The expression level of *FSTOMAGEN* is shown in (Zhao et al. 2022). The results of the statistical test for Fig C and Fig D are in supplementary file. Student t test (unpaired, two-tail) was performed for the Fig B. * represents P<0.05, and n.s. represents no significant difference in statistics.

Three level in biomass was apparently observed (Fig 1 B), and stomatal density gradually increased. Therefore, the lines were divided into three groups to analyze the difference (Fig 1B). The results showed that there was a significant difference in biomass between lower level and intermediate level, or intermediate level and higher level, whether dry weight or fresh weight (Fig 1 B). There was no significant difference in biomass between lower level and higher level (Fig 1 B).

### 2. There was a positive correlation between biomass and water content in different transgenic lines of *Arabidopsis*

In the late stage of growth, the rosette of *Arabidopsis* in some lines turned to purple, brown and dried-up (Supplementary figure 1). Since these transgenic lines had increased stomatal density, the change in rosette might be caused by the excessive loss of water. Additionally, photosynthesis could be significantly affected by the water content of plants. Therefore, we tested the water content of these transgenic plants. We found indeed that the lines with increased stomatal density had lower water content (Fig 2A, supplementary file). Especially, the water content of the 30 lines accompanied with overmuch stomatal density was extremely low (Fig 2A), and the purple and brown of the leaf for this line were most apparent (Supplementary figure 1). More important, the leaf of this line became withered. These results indicated that these transgenic lines experienced water limitation during the growth.

**Figure 2.**
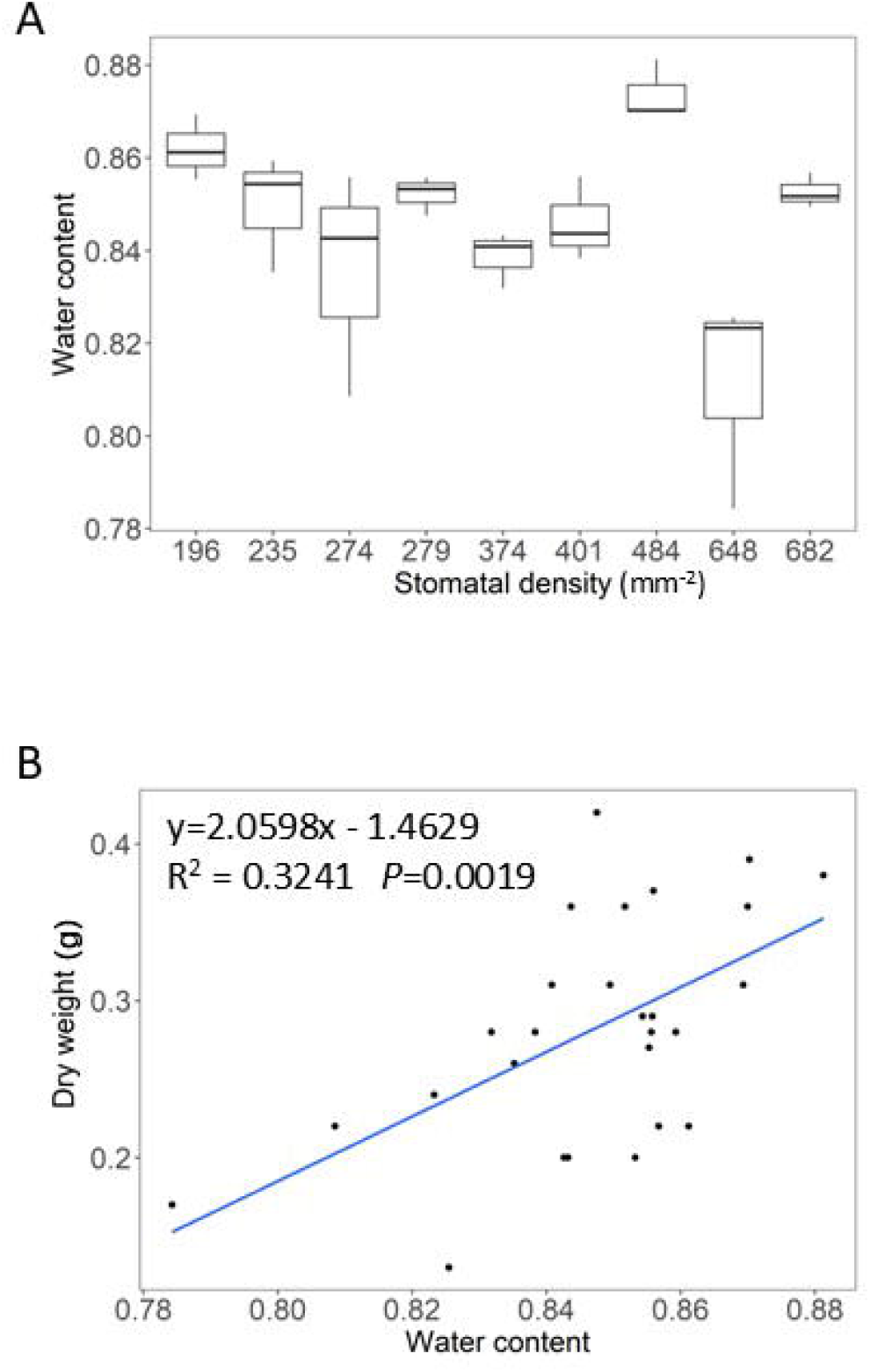
The changes in water content between different *Arabidopsis* lines with the *FSTOMAGEN*overexpression. A, The changes in water content between different *Arabidopsis* lines with different stomatal density. The different stomatal density was achieved by the overexpression of *FSTOMAGEN*. The statistical test is in supplementary file. B, The relationship between dry weight and water content is shown.

The decreased water content in plants probably decrease the photosynthetic rate, and then decrease the biomass of plants. Therefore, we tested the relation between biomass and the water content of plants. Results showed that there was a positive relationship between dry weight and water content (y=−0.8444+1.2575x, R^2^=0.3038,P=0.0052) (Fig 2B).

### 3. The leaf area and leaf growth rate of the transgenic lines with increased stomatal density were increased in the earlier period

There was a large difference in leaf area and leaf growth rate between these lines (Fig 3 A, B, C). The data of the leaf growth was fitted by a growth model, which was approximately an exponential function (Fig 3 D). The leaf growth in the early phase was slower as compared with the leaf growth in later phase (Fig 3 D).We observed the leaf growth rate of these transgenic lines during the earlier period. We found both the leaf area and leaf growth rate of these transgenic lines increased stepwise as stomatal density increase during this period (Fig 3A, B). Especially, because of the stepwise increase, this result accorded to the theory that increased stomatal density induce increased photosynthetic rate. It is worth to note that the pattern of growth rate after 20 days was not similar to the period before this day (Fig 3B).

**Figure 3.**
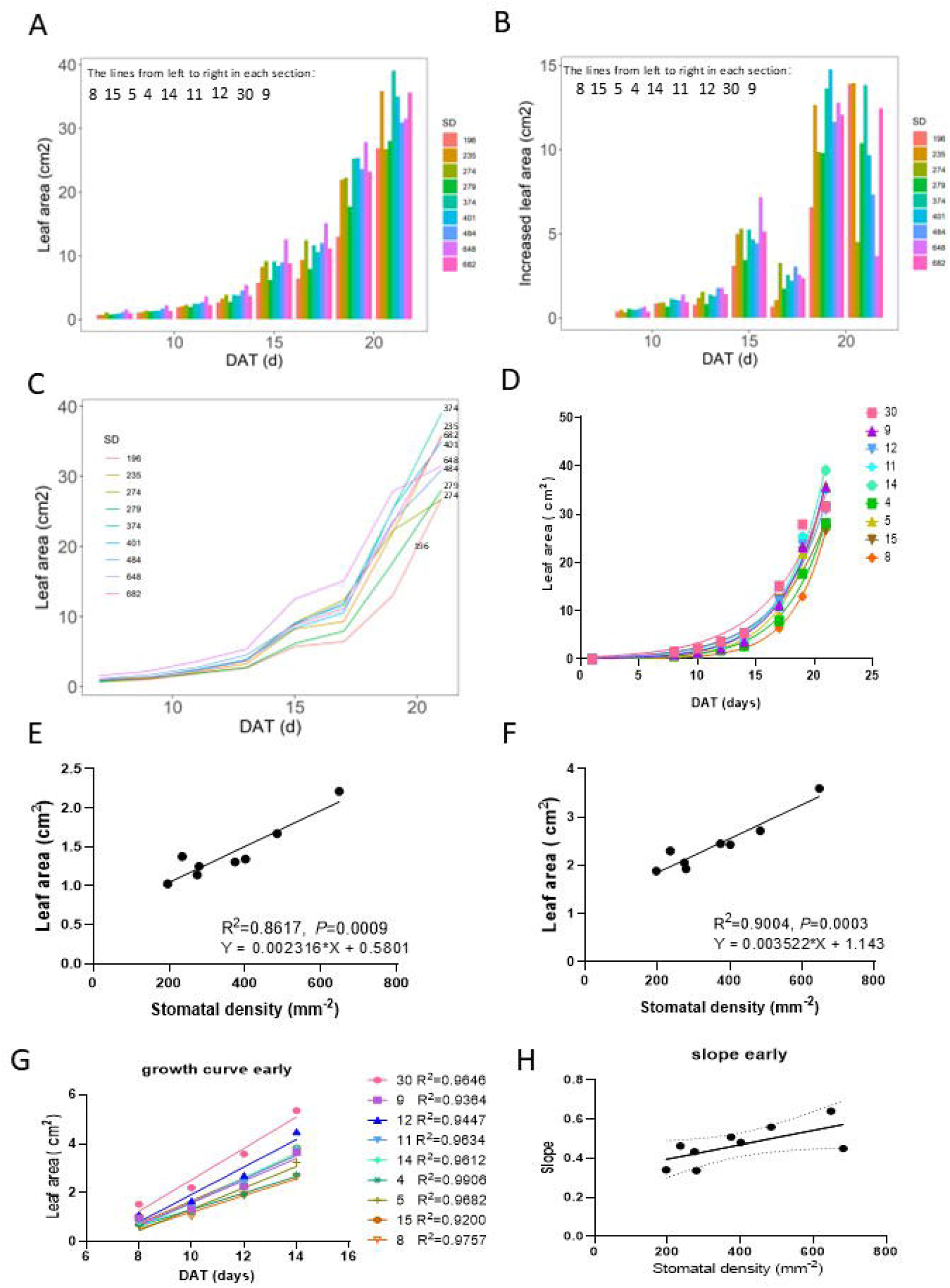
The changes in leaf area between different *Arabidopsis* lines with the *FSTOMAGEN* overexpression at the different date. A, The changes in leaf area between different *Arabidopsis* lines with different stomatal density at the different date. B, The changes in increased leaf area between different *Arabidopsis* lines with different stomatal density at the different date. C, The changed trend in leaf area of different *Arabidopsis* lines during the growth period. The different stomatal density was achieved by the overexpression of *FSTOMAGEN*. D, The fitted growth curve of the growth of leaf area in different trangenic lines. E, The relationship between stomatal density and leaf area at 7-24 without the 682 mm^-2^ stomatal density. F, The relationship between stomatal density and leaf area at 7-26 without the 682 mm^-2^ stomatal density. G, The linear regression between the days and leaf area in the early phase of growth for the different lines. H, The relationship between stomatal density and slope (leaf growth rate in the early phase).

The early phase of leaf growth exhibited apparently linear growth ( Fig 3 D, 3 G). There was a significant linear relationship between the slope of the early phase of leaf growth and stomatal density (Fig 3 H). The leaf growth rate (slope) is strongly and positively correlated to photosynthetic rate, therefore the significant linear relationship between the slope and stomatal density was in line with the notion that increased stomatal density increased photosynthetic rate and furthermore biomass accumulated rate (leaf growth rate almost equal biomass accumulated rate), and the increased biomass accumulated rate caused the final difference in the biomass and leaf area (Fig 1, 3), although this case occurred at the early stage (Fig 3H). There was a significant linear relationships between leaf area and stomatal density at the earlier dates (7/22, 7/24, 7/26 and 7/28) (Supplementary fig 2). We observed a apparent value which likely severally deviated the trend of others (Supplementary fig 2). In order to examine whether it is the real value which severally deviates the trend of others, we performed Internal studentized residual and External studentized residual. The results showed that there indeed was a deviated value (Stomatal density is 682) (Supplementary fig 3). Addtionally, excessive stomatal density may violate the general rule. Therefore, we have sufficient reasons to dislodge this value from the previous regression. After dislodged this value, it was strikingly that there was a strong linear regression between leaf area and stomatal density (Fig 3 E, F), and the R^2^ could reach to about 0.9 (Fig 3 E, F). There was no apparent relationship between stomtatal density and the leaf growth of the later phase (Supplementary fig 3).

## Discussion

### 1) Appropriately increase stomatal density could increase plant biomass

In this study, we focused on the increase in stomatal density by genetically manipulating the expression of the gene belonging to *Flaveria*(the homolog of *STOMAGEN*, the homolog is from *Flaveria*genus). Our results showed that the lines with moderately increased stomatal density had higher biomass as compared to the lines with lower stomatal density (Fig 2A, B). This also indicated that the photosynthetic rate and light use efficiency increased under our growth condition (high light intensity and sufficient water). Our results showed that there was a strong and significant linear relationship between leaf area (or the slope of the leaf area growth) and stomatal density in the earlier phase of *Arabidopsis* growth (Fig 3 E, F, G, H), indicating the increased biomass was caused by the increased stomatal density, and the increased biomass was caused by the accumulation in biomass in the earlier phase of growth.

In agreement with other work(Tanaka et al. 2013), our work also found that the biomass of the line with extremely increased stomatal density didn’t change, and certain line even decreased (Fig 2A, B). The reason for this phenomenon might be that the overmuch stomata largely enhance transpiration, and the amount of the lost water largely exceed the amount of the absorbed water through root. This caused a consequence that plants were at the status with an extreme lack of water (Fig 3A). The extreme lack of water would cause the two occasions which restrict photosynthesis: 1, inducing the decrease in stomatal conductance; 2, weaken the biochemical reaction in photosynthesis. Therefore, the growth of the plants was restricted and the biomass was decreased. The notion that the decreased biomass was caused by the decreased water content in this study was supported by (Fig 3B). That the positive correlation between biomass and water content suggested that decrease in water content weakened the growth of plants and the accumulation of biomass (Figure 3B). Another reason for the decreased biomass in the lines with extremely increased stomatal density should be that energy was overly consumed in the stomatal development and movement. The overly consumption in energy reduced the biomass which should increased in the lines with overmuch stomata.

Our work has shown that the substitution of signal peptide did not affect the function that regulates stomatal density(Zhao et al. 2022), indicating the combination of the functional region and the signal peptide does not affect stomatal density. The function of the functional region is independent from the function of the signal peptide. The function of the signal peptide should be restricted to the accurate location of the gene. Therefore, many types of signal peptides that can accurately locate the gene may be able to replace the own signal peptide of the gene.

### 2) Stomata have large potentials to become the crucial model for the age of quantitative biology in cell level

Genetic engineering in biological science has long been a hotspot. However, many studies focus on the quality in the biological process. Overly changes in the biological process would cause the consequence that is not expected. The results for the lines with overmuch stomata in this study can be an example for this claim (Fig 2A, B).

Our study was an initiating work in the research area where researchers study on the quantity of cells to regulate plants morphology by genetically engineering. In particular, it was the first time that the study showed that the lines with moderately increased stomatal density indeed were able to possess the expected biological process, i.e. increased biomass. There are some limitations for this study. In this study, we used the 35S promoter to launch the overexpression of *FSTOMAGEN*, and selected the lines with different expression. This will encounter multiple troubles. For example, engineering in this study introduced the exogenous gene into the plants. Second, selecting the lines with moderate expression consumed much time. Recently, it is encouraging that a large number of biological techniques has been developed. Among these approaches, the representative widely known is the crispr-cas9(Xing et al. 2014). Directly editing the genome to produce the various expression of the gene might be a better strategy in future.

The molecular mechanism of stomatal development has been well studied. Besides, stomata are convenient to observe and count, and many software tools for counting stomata has been developed in recent years(Fetter et al. 2019)(Wang et al. 2024). More crucially, reduced atmospheric CO_2_ and increased biomass could enable us to attach importance to this work. Therefore, stomata could be the crucial model for the quantitative biology in cell scale.

### 3. Increasing stomatal density to a moderate level through genetically engineering can benefit the reduction of the concentration of atmospheric CO_2_ and production

The genes in regulating stomatal development are functionally conserved across different species (Yin et al. 2017; Lu et al. 2019; Wang et al. 2016; Caine et al. 2018; Hughes et al. 2017). This means that we can modify stomatal density in these species through genetically manipulating the homologs of the genes for regulating stomatal development, which are similar to the experiment conducted in this study.

The increase in the concentration of atmospheric CO_2_ could raise temperature, and this could affect and change global environment(Tonn 2007). The accelerated increase in the concentration of atmospheric CO_2_ could accelerate the change in global climate and environment. This may cause the extreme and rare climate and environment (including the natural hazard like drought and flood). Therefore, that exploring the approach to handle the increase in the concentration of atmospheric CO_2_ is necessary. This study provided insights into the approach. Our study indicated that moderately increased stomatal density can improve the biomass of plants (Fig 1), denoting that increased stomatal density can increase net CO_2_ assimilation rate and enhance the capacity of plants to absorb CO_2_ from external environment. Because the function of genes in regulating stomatal development is conserved between species(Hara et al. 2007; Tanaka et al. 2013; Wang et al. 2016; Karavolias et al. 2023; Shahbaz et al. 2025), and these species play an important role in ecology. It is likely that increasing stomatal density to a moderate level in these species could significantly improve the biomass and benefit the decrease in the concentration of atmospheric CO_2_. Especially, different from C_3_ photosynthesis in *Arabidopsis*, a large number of plants have C_4_ photosynthesis. It is very attractive for us to moderately increase stomatal density to improve biomass by genetic manipulation in C_4_ plants, since C_4_ plants intrinsically have high photosynthetic rate and high production of biomass. Besides, the increased biomass of plants can be used for human. For example, in the industrial and manufacturing sectors, the increased biomass can be used to produce some productions that need cellulose.

### 4) An example for the combination between genetic engineering and the controlling of environment

The traits or characteristics of species enable plants to adapt to certain environment, and this is done by long-term evolution. But in current age, the changes in environment become faster. A faster strategy is required to meet the current changes. Our study suggested that these goals could be reached by manipulating in the molecular levels. The increased stomatal density enables plants to fit the environment with sufficient water and high light intensity. In addition to adapt to natural environments, in future, the environments (sufficient water and high light intensity) in the intelligent plant factory will be suitable to the increased stomatal density. In theory, the characteristics are more complex (For example, biomass), it will be more difficult to enable plants having the expected characteristics through the modification from the single gene in genetic engineering. Additionally, most current goal of the genetic engineering is to find the genes that function in most, changed and even all environments. This seems not realistic. This largely restricts the application and the development of genetic engineering and molecular experiment. It is believed that most genes have their roles, and when in corresponding environment, they can exhibit their function in regulating the traits. Our study proposed that the cooperation between the genetic manipulation and the controlling of environment could solve this trouble. In this study, the biomass of *Arabidopsis* increased under a higher light intensity and sufficient water. Although we didn’t perform the experiment under lower light intensity, increasing stomatal density should not improve the biomass since the photosynthetic biochemistry is limited. Therefore, this study is a valuable example and provides the new insights for these kinds of studies.

## Acknowledgement

Thanks for the cooperating of the experiments with Fu sang Liu. Thanks for the cooperating from the groups members.

## Author contribution

Yong-yao Zhao designed the research in this article and wrote the article. Yong-yao Zhao leaded and performed the experiment. Yong-yao Zhao analyzed the data. Yong-yao Zhao conducted the graphing and plotting.

## Figure Legends

Suppmentary figure 1 The photos at the different date during the growth for the different *Arabidopsis* lines with the *FSTOMAGEN*overexpression.

The photos for the different *Arabidopsis* lines with different stomatal density at 19 day (A), 22 day (B), 30 day (C), 33 day (D). Stomatal density is shown above the photos.

Supplementary figure 2 The relationship between leaf area and stomatal density.

The relationship between leaf area and stomatal density at 2022-7-22 (A), 2022-7-24(B), 2022-7-26(C), 2022-7-28(D), 2022-8-01(E), 2022-8-03(F), 2022-8-05(G).

Supplementary figure 3 The leaf growth rate in the later growth phase

A, The linear regression between the days and leaf area in the later phase of growth for the different lines. B, The relationship between leaf growth rate (slope) and stomatal density.

Supplementary table 1 The internal studentized residual and the external studentized residual for the relationship between leaf area and stomatal density.

Supplementary file The statistic test for the Figure 2 and 3.

## Notes

### Competing Interest Statement

The authors have declared no competing interest.

### Summary of Updates

Add the figure and revise the sentence in the paper

