## Supplementary figures and images for "Quantifying cells : moderately increasing stomatal density through overexpressing *FSTOMAGEN* improved the biomass of *Arabidopsis*"

### Supplementary figure 1

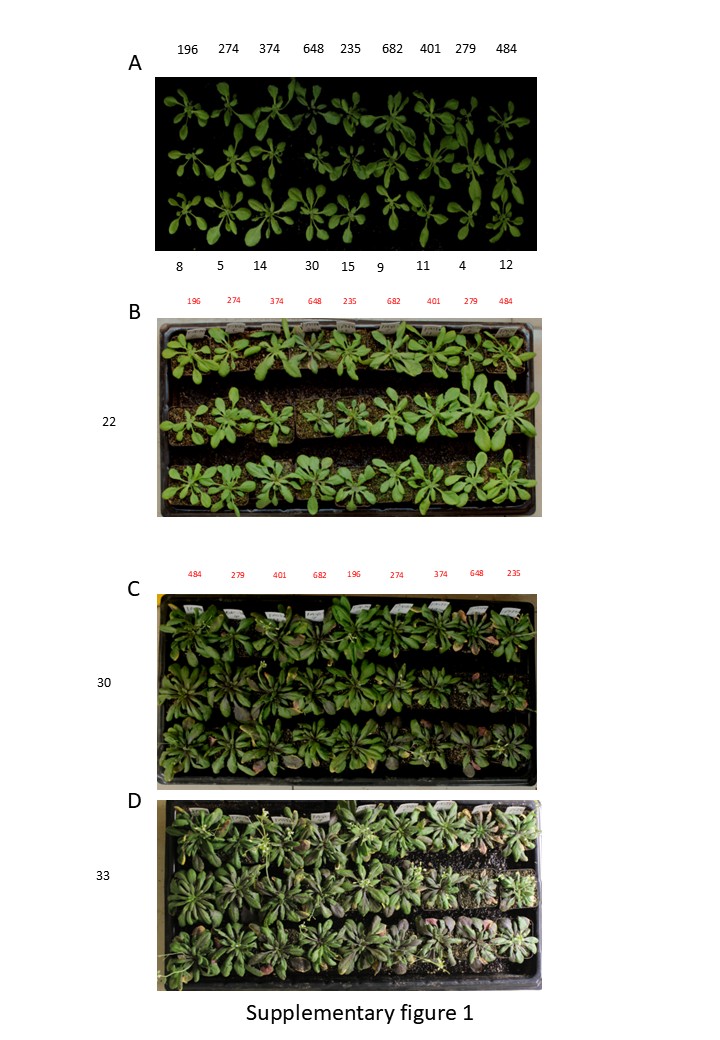

### Supplementary figure 2

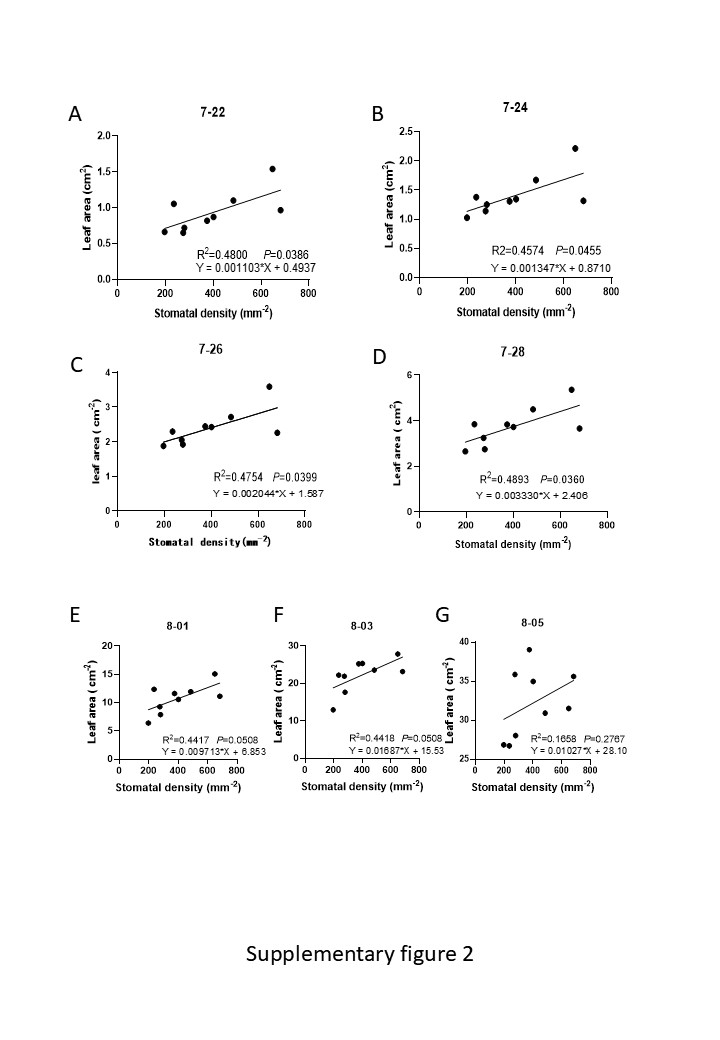

### Supplementary figure 3

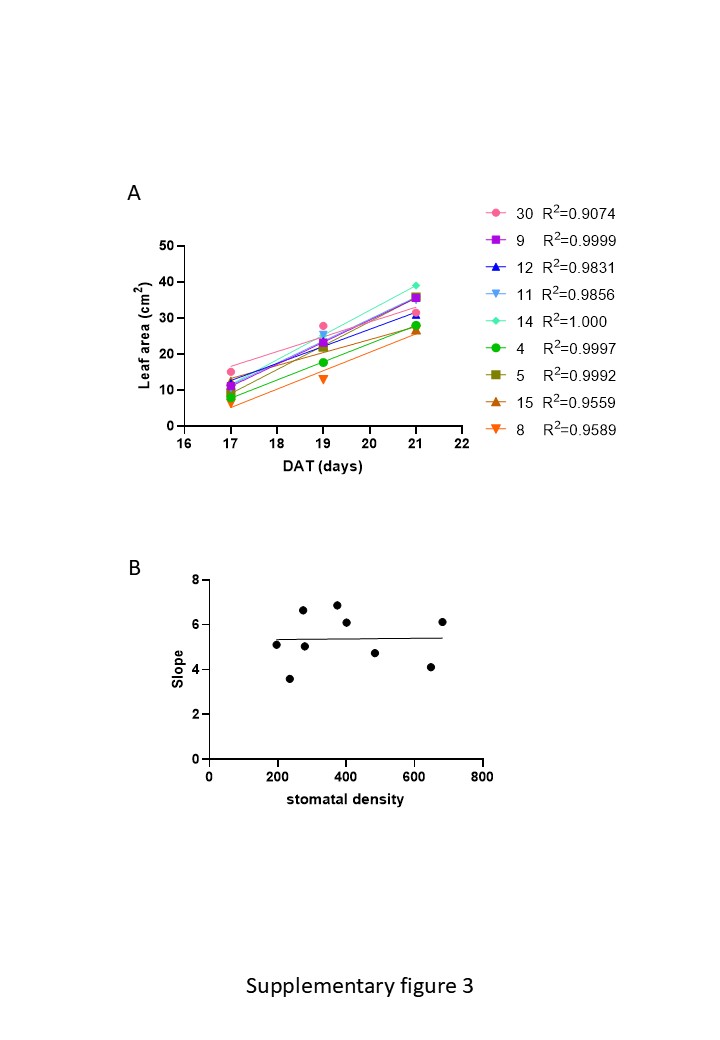

### Supplementary table 1

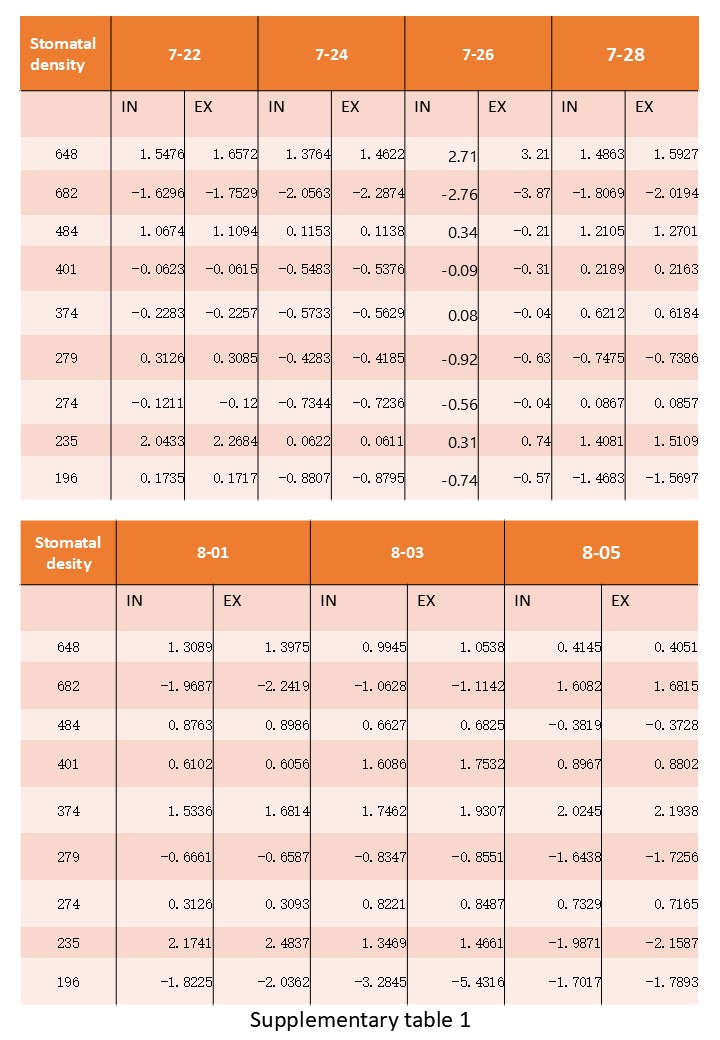
